# Ancestral chromatin configuration constrains chromatin evolution on differentiating sex chromosomes in *Drosophila*

**DOI:** 10.1101/019786

**Authors:** Qi Zhou, Doris Bachtrog

## Abstract

Sex chromosomes evolve distinctive types of chromatin from a pair of ancestral autosomes that are usually euchromatic. In Drosophila, the dosage-compensated X becomes enriched for hyperactive chromatin in males (mediated by H4K16ac), while the Y chromosome acquires silencing heterochromatin (enriched for H3K9me2/3). Drosophila autosomes are typically mostly euchromatic but the small dot chromosome has evolved a heterochromatin-like milieu (enriched for H3K9me2/3) that permits the normal expression of dot-linked genes, but which is different from typical pericentric heterochromatin. In *Drosophila busckii*, the dot chromosomes have fused to the ancestral sex chromosomes, creating a pair of ‘neo-sex’ chromosomes. Here we collect genomic, transcriptomic and epigenomic data from *D. busckii*, to investigate the evolutionary trajectory of sex chromosomes from a largely heterochromatic ancestor. We show that the neo-sex chromosomes formed <1 million years ago, but nearly 60% of neo-Y linked genes have already become non-functional. Expression levels are generally lower for the neo-Y alleles relative to their neo-X homologs, and the silencing heterochromatin mark H3K9me2, but not H3K9me3, is significantly enriched on silenced neo-Y genes. Despite rampant neo-Y degeneration, we find that the neo-X is deficient for the canonical histone modification mark of dosage compensation (H4K16ac), relative to autosomes or the compensated ancestral X chromosome, possibly reflecting constraints imposed on evolving hyperactive chromatin in an originally heterochromatic environment. Yet, neo-X genes are transcriptionally more active in males, relative to females, suggesting the evolution of incipient dosage compensation on the neo-X. Our data show that Y degeneration proceeds quickly after sex chromosomes become established through genomic and epigenetic changes, and are consistent with the idea that the evolution of sex-linked chromatin is influenced by its ancestral configuration.

**Author Summary:** DNA is packaged with proteins into two general types of chromatin: the transcriptionally active euchromatin and repressive heterochromatin. Sex chromosomes typically evolve from a pair of euchromatic autosomes. The Y chromosome of Drosophila is gene poor and almost entirely heterochromatic; the X chromosome, in contrast, has evolved a hyperactive euchromatin structure and globally up-regulates its gene expression, to compensate for loss of activity from the homologous genes on the Y chromosome. The evolutionary trajectory along which sex chromosomes evolve such opposite types of chromatin configurations remains unclear, as most sex chromosomes are ancient and no longer contain signatures of their transitions. Here we investigate a pair of unusual young sex chromosomes (termed ‘neo-Y’ and ‘neo-X’ chromosomes) in *D. busckii,* which formed through fusions of a largely heterochromatic autosome (the ‘dot chromosome’) to the ancestral sex chromosomes. We show that nearly 60% of the neo-Y genes have already become non-functional within only 1 million years of evolution. Gene expression is lower on the neo-Y than on the neo-X, which is associated with a higher level of binding of a silencing heterochromatin mark. The neo-X, on the other hand, shows no evidence of evolving hyperactive chromatin for dosage compensation. Our results show that the Y chromosome can degenerate quickly, but the tempo and mode of chromatin evolution on the sex chromosomes may be constrained by the ancestral chromatin configuration.

## Introduction

Sex chromosomes have originated independently many times from ordinary autosomes in both plants and animals [1]. A common feature of heteromorphic sex chromosomes is that while X chromosomes maintain most of their ancestral genes, Y chromosomes often degenerate due to their lack of recombination, with only few functional genes remaining (for a recent review see [2]). The loss of gene function is often accompanied by an accumulation of repetitive DNA on ancient Y chromosomes, and a switch of chromatin structure from euchromatin to genetically inert heterochromatin [2, 3]. Loss and silencing of Y-linked genes drives the evolution of dosage compensation on the X chromosome, which is often mediated by chromosome-wide epigenetic modifications. Drosophila males, for example, acquire a hyperactive chromatin conformation of their single X, while one of the two X’s in female mammals becomes heterochromatic [4, 5].

Studies of young sex chromosomes have improved our understanding of the genomic and epigenomic mechanisms driving the divergence between X and Y [6-9]. Neo-sex chromosomes of Drosophila are formed by chromosomal fusions between the ancestral sex chromosomes and ordinary autosomes. The neo-Y, which is the autosome that became linked to the Y, entirely lacks recombination since it is transmitted through males only, which in Drosophila do not undergo meiotic recombination. Consistent with theoretical predictions that selection is ineffective on non-recombining chromosomes [10], neo-Y chromosomes in several Drosophila taxa have undergone chromosome-wide degeneration, and the extent of gene loss roughly corresponds to the age of the neo-Y. In particular, the very recently formed neo-Y of *D. albomicans* (<0.1 million year old) still contains most of its protein coding genes with <2% being putatively non-functional [11], but a large fraction of neo-Y genes (roughly 40%) are down-regulated [9], suggesting that transcriptional silencing might be initiating Y degeneration. The older neo-Y chromosome of *D. miranda* (1.5 million years old) has acquired stop codons and frame-shift mutations in almost half of its genes, shows a dramatic accumulation of transposable elements (between 30-50% of its DNA is composed of TEs) [12, 13], and most neo-Y genes are expressed at a lower level than their neo-X homologs [11]. These changes at the DNA sequence level are accompanied by a global change in chromatin structure, and the *D. miranda* neo-Y is adopting a heterochromatic appearance marked by histone H3 lysine 9 di-methylation (H3K9me2) [3]. The neo-X of *D. miranda*, in contrast, has maintained most of its ancestral genes but is evolving partial dosage compensation, by co-opting the canonical dosage-compensation machinery of Drosophila (the *MSL*-complex). This complex is targeted to the ancestral X of Drosophila species, and up-regulates gene expression through changes of the chromatin conformation of the X, mediated by histone H4 lysine 16 acetylation (H4K16ac) [14, 15]. The neo-sex chromosome shared by members of the *D. pseudoobscura* species group was formed about 15 million years ago, and has evolved the typical properties of old sex chromosomes: the neo-Y is completely degenerate and heterochromatic, while the neo-X is fully dosage compensated by the *MSL* machinery [3, 16].

Well-studied neo-sex chromosome systems are all derived from euchromatic autosomes, and studying a neo-sex chromosome that originated from an autosome with some features similar to heterochromatin may allow a more general understanding of the evolutionary principles of chromatin formation on sex chromosomes. Here, we collect data on the genome, transcriptome and epigenome of *D. busckii*, a species with a poorly characterized neo-sex chromosome derived by a fusion (and supposedly followed by a pericentric inversion on the X) between the ancestral sex chromosomes and the “heterochromatic*”* dot chromosome (**Figure 1A**) [17, 18]. The age, and the extent of sequence, expression and epigenetic divergence of the neo-sex chromosomes of *D. busckii* are unknown, but the dot chromosome has an unusual evolutionary history and a unique chromatin structure. It was a sex chromosome in an ancestor of higher Diptera, and only reverted to an autosomal inheritance in the ancestor of the Drosophilidae family [19, 20]. Studies on the assembled distal arm (∼1.2Mb) of the *D. melanogaster* dot chromosome have revealed several features that distinguish it from other autosomes: it has a very low recombination rate and a high repeat content [21-23], harbors less than 100 genes [24] that have low codon usage bias[25] and which show evidence of reduced levels of positive and purifying selection [26]. Genes on the dot chromosome are embedded into a unique heterochromatin-like milieu that is regulated differently from canonical pericentric heterochromatin [21, 27]. Both dot-linked genes and genes located in pericentric heterochromatin are enriched for the ‘silencing’ histone marks H3K9me2 and H3K9me3 and the heterochromatin protein *HP1a* relative to euchromatin, but show a depletion of these marks at the transcriptional start sites of active genes. In addition, expression of dot-linked genes (but not genes in pericentric heterochromatin) requires binding of the chromosomal protein *Painting of Fourth* (*POF*) and the histone methyltransferase *EGG*, and the gene bodies of transcribed genes show an enrichment of the histone modification H3K9me3 (but not H3K9me2) that is not observed at active genes located in pericentromeric heterochromatin. Genes on the dot chromosome that are not expressed and repetitive regions on the dot chromosome probably adopt a more general *POF*/*EGG* independent mechanism of heterochromatin packaging that is shared with pericentromeric regions [28].

**Figure 1.**
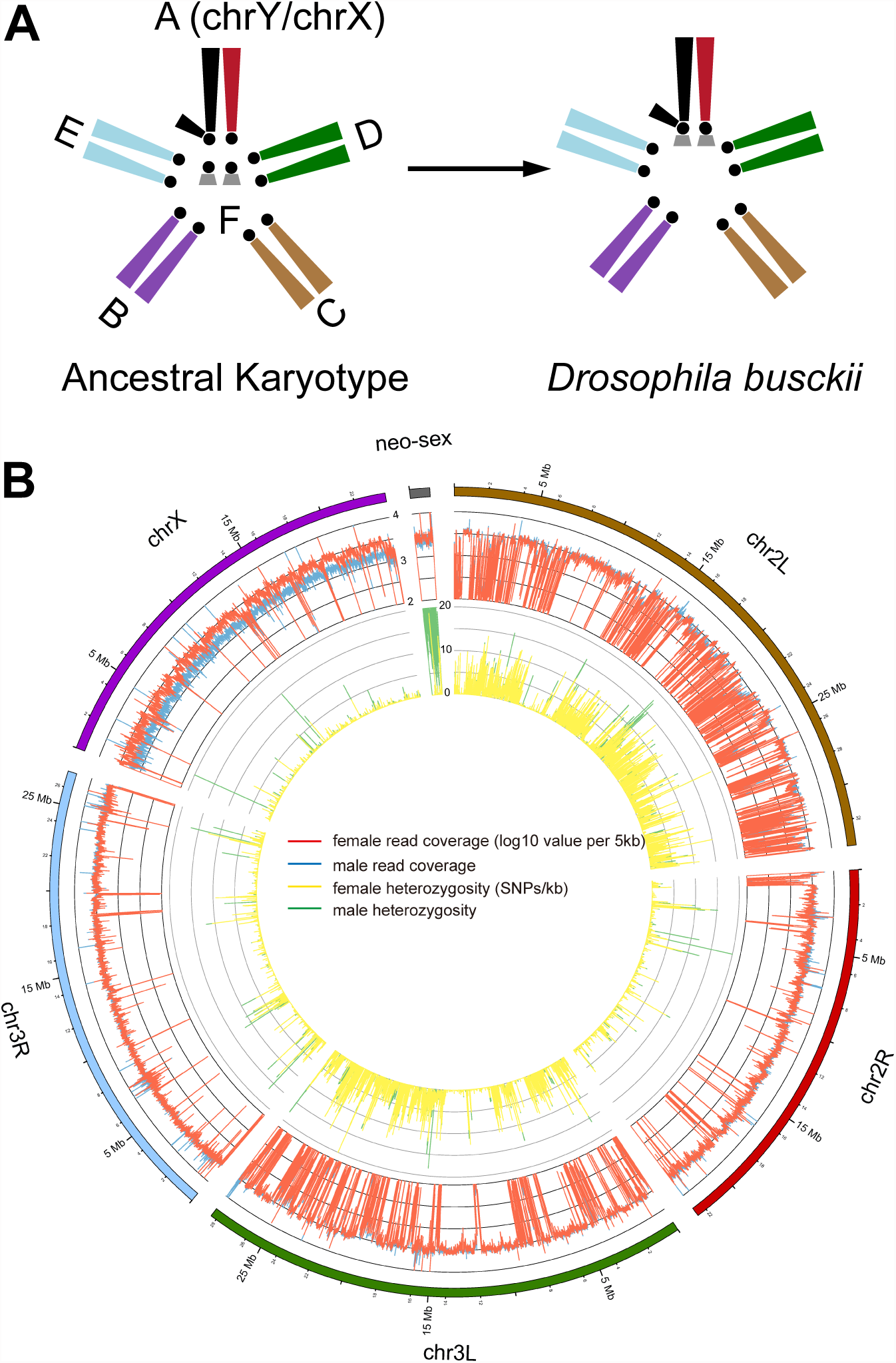
Karyotype and genome of *D. busckii*. **A.** Karyotype of *D. busckii*. The ancestral karyotype of Drosophila species consists of six chromosomal arms termed ‘Muller’s elements’. Muller-A element is the ancestral sex chromosome shared by all Drosophila species, and Muller-F is the dot chromosome. In *D. busckii,* the dot chromosome pair has fused to the ancestral X and Y chromosome and became the neo-X and neo-Y chromosome. **B.** Coverage and heterozygosity patterns of the *D. busckii* genome. For each chromosome of *D. busckii* (named after its homologous chromosome in *D. melanogaster*), we show mapped read coverage in male (blue) and female (red), and single nucleotide polymorphism (SNP) density (sites/kb) within 5kb non-overlapping windows along the chromosome. chrX shows a reduction of male coverage because the ancestral Y chromosome is completely degenerated in male *D. busckii*. The neo-sex (dot) chromosome shows similar coverage to autosomes, indicating that most neo-Y reads can still be mapped to the neo-X. The increase of male SNP density on the neo-sex chromosome indicates sequence divergence between the neo-X and neo-Y alleles.

Intriguingly, in three Drosophila species including *D. busckii*, *POF* was found to bind the X chromosome specifically in males [29]. This mimics the localization of the *MSL* complex, the canonical dosage compensation machinery of Drosophila, but unlike in other Drosophila species, immunostaining to polytene chromosomes detected no binding of the *MSL* complex on the X chromosome of *D. busckii* [16, 29]. The phylogenetic position of *D. busckii* is uncertain, and some early studies placed it as a sister to all other Drosophila species [16, 30]. These findings, together with the discovery that the dot was actually the ancestral sex chromosome in Diptera led to the hypothesis that *D. busckii* might harbor a more ancestral mechanism of dosage compensation mediated by *POF*, which may have been derived from a dosage compensation system in an ancestor of Drosophilidae where the dot was the X chromosome [19]. Here, we collect DNA sequence, transcriptome and chromatin data characteristic of dosage compensation and heterochromatin together with immunostaining of polytene chromosomes, to characterize the formation of a sex chromosome from a heterochromatic ancestor, and also to disentangle the relationship between *POF* and *MSL*.

## Results

### Genome assembly and annotation

We sequenced the *D. busckii* female genome to an extremely high sequencing coverage (>150 fold, **Table S1**) with libraries spanning a gradient of insert sizes (up to 10kb) to produce a highly continuous *de novo* assembly (scaffold N50: 946kb, average scaffold size: 60.8kb) with a total assembled length of 152.7Mb. Orthologous Drosophila chromosomes show high conservation (>95%) in their gene content [32], and we assign the chromosomal locations of *D. busckii* genome scaffolds based on their alignments with *D. melanogaster* chromosomes. 89% of the sequences could be assigned to individual linkage groups, and we further tested our chromosomal assignments by sequencing the male genome. The ancestral X chromosome is hemizygous in males, and mapped male read depth is indeed only half of the female read depth along the entire X chromosome, while read depths are very similar between sexes on autosomes (median log10 coverage value of male vs. female: 3.50 vs. 3.46; *P*>0.05, Wilcoxon test) (**Figure 1B**). Interestingly, coverage in both sexes is also very similar along the dot chromosome and only slightly reduced in males (median of male vs. female: 3.42 vs. 3.44), implying that the neo-X and neo-Y still share considerable sequence homology. This suggests that the age of the neo-sex system of *D. busckii* is younger than that of *D. miranda*, which shows significantly reduced male read depth (by about 25%) along its neo-sex chromosome due to neo-X/Y divergence [11].

We annotate 13.1% of the assembled genome as consisting of repetitive elements, with the dot chromosome containing the highest repeat content (17.3%) among all chromosomes. We also produce transcriptomes of male and female *D. busckii* third instar larvae and adults, and integrated them during gene annotation. A total of 12,648 protein-coding genes were annotated using *D. melanogaster* proteins as query, 11,859 (93.6%) of which have one-to-one *D. melanogaster* orthologs. We find a higher proportion of annotated genes actively expressed in male than in female (69.4% vs. 53.8%) with a normalized expression level RPKM (average RNA-seq reads per kilobase of gene per million fragments mapped) higher than 5, and also a generally lower male expression level on the X chromosome relative to autosomes (Wilcoxon test, *P*<0.05, **Figure S1**), in both developmental stages. These patterns are consistent with sex-biased expression patterns found in *D. melanogaster* [33, 34], and a similar ‘demasculinization’ found on the X chromosomes in other Diptera [19, 20].

### Phylogenetic position of *D. busckii*

The phylogenetic relationship of *D. busckii* within the Drosophila genus is unclear. Some studies placed it as a sister to all other Drosophila species [16, 30], while others put it within the Drosophila subgenus [35]. This uncertainty in the phylogenetic position of *D. busckii* could have resulted from the small number of genes that were previously investigated, and we use whole-genome sequence alignments of representative Drosophila species and other Drosophilidae, to generate a phylogenomic tree. Our alignments include *D. melanogaster*, *D. pseudoobscura* and *D. willistoni* from the Sophophora subgenus; *D. albomicans* [11], *D. grimshawi* and *D. virilis* from the Drosophila subgenus, *D. busckii* and two recently sequenced Diptera species within the Drosophilidae family: *Scaptodrosophila lebanonensis* [36] and *Phortica variegata* [19] as outgroups to the Drosophila genus [35-37]. In total, we aligned CDS sequences of 6189 orthologous genes spanning a total of 19.1Mb from each species and acquired a consensus tree with high bootstrapping values (**Figure 2**). *D. busckii* consistently clusters with the Drosophila subgenus species (*D. albomicans*, *D. grimshawi* and *D. virilis*) rather than being placed at the base of all Drosophila. This phylogenetic analysis suggests that *D. busckii* is not a member of an early divergent Drosophila lineage, but originated within the Drosophila subgenus.

**Figure 2.**
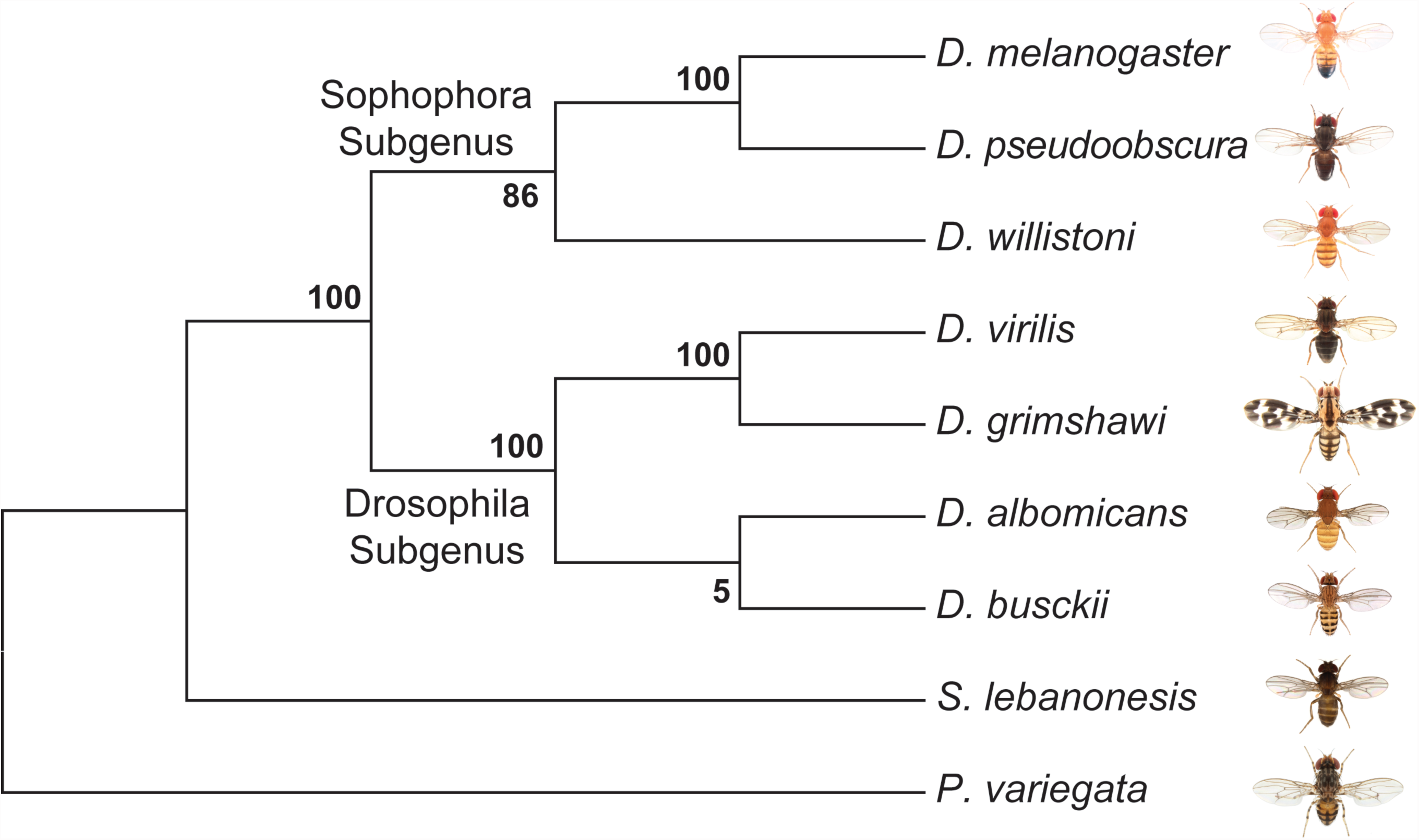
Phylogeny of *D. busckii*. We used 6189 orthologous gene pairs from 9 Diptera species and constructed a phylogenomic tree. Although the bootstrap value at the ancestral node of *D. busckii* and *D. albomicans* is low, *D. busckii* is grouped with high confidence within the Drosophila subgroup instead of as a sister group to all Drosophila species, as previously hypothesized [16, 30].

### Sequence degeneration of the *D. busckii* neo-Y

We assembled and mapped a total of 1.17Mb (with 6.9% of the sequence as gaps) of dot chromosome sequence in *D. busckii*, in comparison to 1.35Mb of assembled dot sequence in *D. melanogaster*. The *D. busckii* dot chromosome overall shows more than 10 times higher levels of heterozygosity (1.56 SNPs per 100bp on average) in male than in female, predominantly due to nucleotide sequence divergence between the neo-X and neo-Y chromosomes (**Figure 1B**). The median level of pairwise divergence at silent sites between neo-X and neo-Y alleles is 0.84%, which is about 3 times lower than synonymous divergence between neo-sex-linked genes of *D. miranda* (2.8%) [11]. Assuming a mutation rate of 5 × 10^-9^ per bp (as estimated in *D. melanogaster*) [38] and 10 generations a year, this indicates that the *D. busckii* neo-sex chromosomes originated only about 850,000 years (0.85 MY) ago. Note that while the fixation of ancestral polymorphisms can contribute to the neo-X/Y divergence, the low level of silent site diversity on the dot [23] implies that ancestral polymorphism is expected to have very limited impact on our estimate of the age of the neo-sex chromosomes of *D. busckii*. The recent formation of the *D. busckii* neo-sex chromosome is consistent with the similar level of read depth observed between sexes along the neo-X chromosome, suggesting this system is still at an initial stage of differentiation (**Figure 1B**).

We annotate a total of 86 neo-sex linked genes (vs. 80 protein-coding genes on the *D. melanogaster* dot chromosome, see notes in Materials and Methods), all of which show the same level of read depth between sexes (**Figure S2**). Thus, unlike on the older neo-Y chromosome of *D. miranda* [11], none of the protein-coding genes has yet been deleted from the neo-Y of *D. busckii*. However, we find male-specific SNPs or indels (i.e., mutations on the neo-Y) that cause premature stop codons and/or frameshift mutations in 50 neo-sex linked genes, implying that there is a large number of genes on the neo-Y that supposedly have lost their normal functions (**Figure 3A**). The proportion of putative non-functional genes (58.2%) is much higher on the neo-Y of *D. busckii* than on that of *D. miranda* (34.2%) [11].

**Figure 3.**
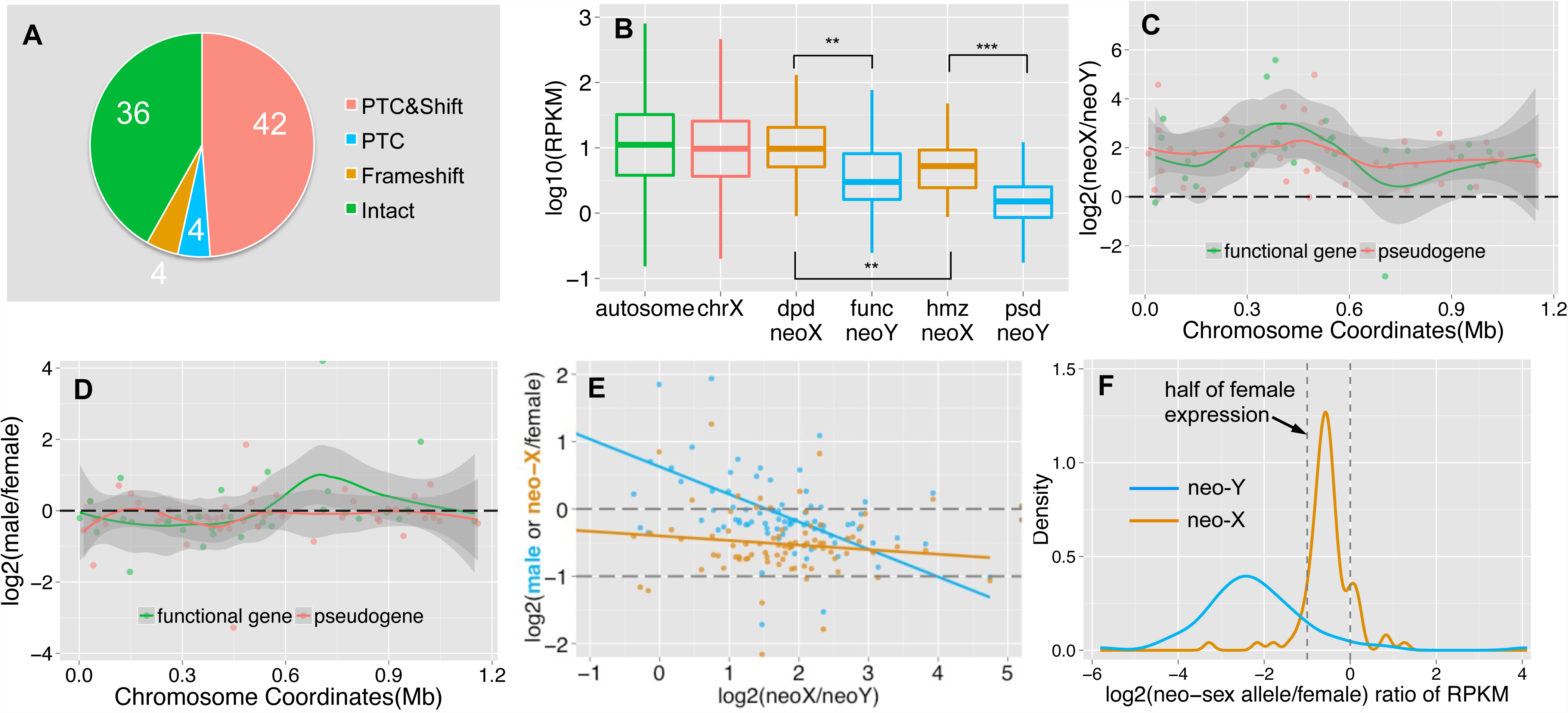
Functional degeneration of neo-Y genes. **A.** Composition of neo-Y linked genes. We show numbers of putative functional genes (‘Intact’), genes with premature stop codons (‘PTC’) and/or frameshift (‘Shift’) mutations on the neo-Y. **B**. Boxplots of gene expression level on each chromosome. We divide neo-sex linked genes according to the functional status of the neo-Y genes: functional (func) neo-Y genes, and their diploid (dpd) neo-X homologs; non-functional (psd) neo-Y genes and their hemizygous (hmz) neo-X homologs. The former group of neo-sex linked genes shows a higher expression level than the latter. **C.** Allelic expression bias of neo-sex linked genes in male adults. Shown are the log ratios of neo-X expression vs. neo-Y expression along the neo-sex chromosome, with putatively functional neo-Y genes in red and pseudogenes in green. We also plot the loess smooth lines separately for the two categories of genes, in order to show the local variation of the log ratio along the chromosome position. Any genes above 0 have higher neo-X expression relative to the neo-Y. **D.** Sex-bias expression of neo-sex linked genes. We show the expression difference between sexes for neo-sex linked genes, with neo-X/Y gene expression level combined in male, and only neo-X gene expression in female. **E.** Correlation between relative neo-sex allelic expression vs. sex-biased expression and relative neo-X expression. Shown are the ratios of neo-X vs. neo-Y expression level for neo-sex linked genes, vs. their expression ratio between sexes (in blue), and the ratio of neo-X expression in male vs. that in female (in orange), as well as their linear regression lines. **F.** Density plot of the ratios of male neo-sex alleles (neo-X in orange, neo-Y in blue) vs. female expression levels. Assuming an equal expression level between sexes, we expect the distribution of relative neo-X alleles’ expression to be around half of the female expression level (dashed line).

This is unexpected, since there has been less time for degeneration on the younger neo-Y chromosome of *D. busckii*. In addition, the much smaller size of the dot chromosome predicts weaker effects of Hill-Robertson interference [10, 39] and thus a lower rate of degeneration on the *D. busckii* neo-Y. However, simulation results have shown that the effects of interference asymptote quite fast with the number of genes [39, 40]. Several other factors could help to explain the large fraction of non-functional genes on the recently formed neo-Y of *D. busckii*. First, genes located on the dot generally show lower levels of evolutionary constraint [41, 42]. Consistent with reduced levels of purifying selection on dot-linked genes, we find that the neo-X alleles show a significantly lower level of codon usage bias than genes on autosomes and the X chromosome (Wilcoxon test, *P*<0.05; **Figure S3**). Note that it is possible that selection for optimal codon usage has become more efficient for dot-linked genes on the neo-X since the dot/X fusion, which may have placed them within a more highly recombining environment, as has been observed for *D. willistoni* [43]. In this case, ancestral levels of codon usage bias may have been even lower for dot-linked genes.

Further, the median rate of protein evolution (as measured by the ratio of nonsynonymous vs. synonymous substitutions using PAML) at the ancestral branch before the neo-X/Y divergence is higher than that of other autosomes (median *K*_a_/*K*_s_ = 0.082 vs. 0.075), and non-functional genes show a higher ancestral rate of protein evolution than genes with a functional copy on the neo-Y (median *K*_a_/*K*_s_ = 0.086 vs. 0.068; **Figure S4**). Although both differences are not statistically significant, probably due to the low number of genes on the dot chromosome, these results are consistent with the idea that genes under lower selective constraints are becoming pseudogenized more quickly on a degenerating neo-Y, as observed on the neo-Y of *D. miranda* [11, 44]. In addition, the gene content of the dot chromosome appears feminized / demasculinized, that is, dot genes in Drosophila and in other Diptera species are over-expressed in ovaries, and under-expressed in testis [19]. Genes with female function are under less purifying selection on the male-limited neo-Y chromosome, which may contribute to accelerated rates of pseudogenization. Neo-X homologs of neo-Y genes that are functional are expressed at a significantly higher level in both male larvae (**Figure S5B**) and adults (**Figure 3B**) than neo-X homologs of neo-Y genes that have become pseudogenized (Wilcoxon test, *P*=0.0087). This indicates that the loss of functional Y-linked genes preferentially starts from lowly-expressed genes with less selective constraints, consistent with our findings on the neo-Y of *D. miranda* [44]. Finally, hemizygosity of dot-linked genes is generally tolerated in *D. melanogaster* [42], and null mutations at dot-linked genes may have a negligible effect on fitness if heterozygous. Thus, lower levels of evolutionary constraints, an excess of female-biased genes, and general haplosufficiency of dot genes may contribute to their rapid degeneration on the neo-Y of *D. busckii*.

### Transcriptomic and Epigenomic Evolution of the neo-Y

In addition to functional decay in protein coding sequences, we also found a chromosome-wide expression bias for neo-sex linked genes (**Figure 3C**): 75 genes (88%) display significantly higher expression from the neo-X chromosome relative to the neo-Y in male adults (Fisher’s exact test, *P*<0.05, see Methods), and a similar pattern was observed in male larvae (**Figure S3A**). Putative pseudogenes on the neo-Y tend to show a slightly more severe (but not statistically significant) expression bias than functional genes (median log2 ratio of neo-X vs. neo-Y expression: 1.80 vs. 1.71; Wilcoxon test, *P*=0.41). This chromosome-wide expression bias for neo-sex linked genes could be caused by down-regulation of neo-Y alleles and/or up-regulation of neo-X alleles (i.e., dosage compensation). Although many genes (77.9%) show a similar level of expression for male (with neo-X/Y gene expression levels combined) and female (less than 1.5 fold difference; **Figure 3D**), genes with lower relative expression from the neo-Y tend to be more female-biased (**Figure 3E**, blue line, Spearman’s rank correlation coefficient: −0.47, *P*=1.04e-5). This suggests that neo-X-biased expression is partly due to down-regulation of neo-Y linked genes. The single neo-X chromosome in males is transcribed at a higher level than a single neo-X chromosome in females (**Figure 3F**), which suggests that some form of dosage compensation has evolved on the neo-X. However, there is no significant correlation between down-regulation of neo-Y genes (i.e. neo-X vs. neo-Y expression bias), and up-regulation of neo-X genes in males (i.e. expression of the neo-X in males vs. females, **Figure 3E**, orange line; *F*-statistic *P*>0.05). This may suggests that dosage compensation is not gene-specific, but could also reflect a lack of statistical power due to the low number of genes on the dot.

The neo-Y chromosome of *D. miranda* has become partially heterochromatic within 1.5 million years. It is enriched for the silencing histone modification H3K9me2 relative to the neo-X and other chromosomes [3], and expression of neo-Y genes is down-regulated chromosome-wide. To investigate whether an accumulation of silencing histone marks may cause down-regulation of neo-Y linked gene expression in *D. busckii*, we obtained ChIP-seq profiles of both H3K9me2 and H3K9me3 from male larvae. The two histone modification marks are strongly correlated with each other and *HP1a* in pericentric heterochromatin, but have distinctive distributions on the dot chromosome of *D. melanogaster*: H3K9me3 shows an unusual correlation with *POF* over actively transcribed gene bodies, while H3K9me2 strongly associates with silenced genes [27, 45].

We analyzed the distribution of H3K9me2 and H3K9me3 at active and silent genes (expression status defined from **Figure S1**), and find that both marks are significantly enriched on the dot chromosomes of *D. busckii* relative to autosomes (Wilcoxon test, *P*<0.05; see Methods, **Figure 4A, D**). H3K9me3 shows a similar level of enrichment between the neo-Y and the neo-X (Wilcoxon test, *P*>0.05, **Figure 4D**), and enrichment tends to be higher at active relative to silent genes on both the neo-X and neo-Y (Wilcoxon test *P>*0.05; **Figure 4D-F**). In contrast, H3K9me2 levels are significantly increased at neo-Y genes relative to their neo-X homologs (Wilcoxon test, *P*=0.000637, **Figure 4A**), particularly on those that are transcriptionally silenced (Wilcoxon test, *P*=0.000381, **Figure 4A-C**), and non-functional neo-Y genes show a significant increase in H3K9me2 binding relative to their neo-X homologs (Wilcoxon test, *P*=0.0001494; **Figure S6**). The H3K9me2 enrichment level of silent neo-Y genes is higher than that of active neo-Y genes (median value: 0.79 vs. 0.47, Wilcoxon test *P*=0.089, **Figure 4A**), and the enrichment level of H3K9me2, but not H3K9me3, is negatively correlated with the gene expression level of neo-Y but not neo-X alleles (**Figure S7,** Spearman’s rank correlation coefficient −0.23, *P*=0.04). We further analyzed metagene enrichment profiles, and find both H3K9me2 and H3K9me3 to be enriched at gene bodies relative to their flanking regions. The increase of H3K9me2 enrichment on silent neo-Y genes is not restricted to gene bodies but extends into flanking regions as well (**Figure 4C**). These results suggest that down-regulation of neo-Y gene expression may be caused by H3K9me2 modification, but it is also possible that some genes are first silenced through mutations in their regulatory region, and then preferentially become targeted by H3K9me2. Overall, our results provide robust evidence that the neo-Y chromosome of *D. busckii* is becoming more heterochromatic, mediated by H3K9me2 enrichment, which further contributes to the degeneration of neo-Y genes.

**Figure 4.**
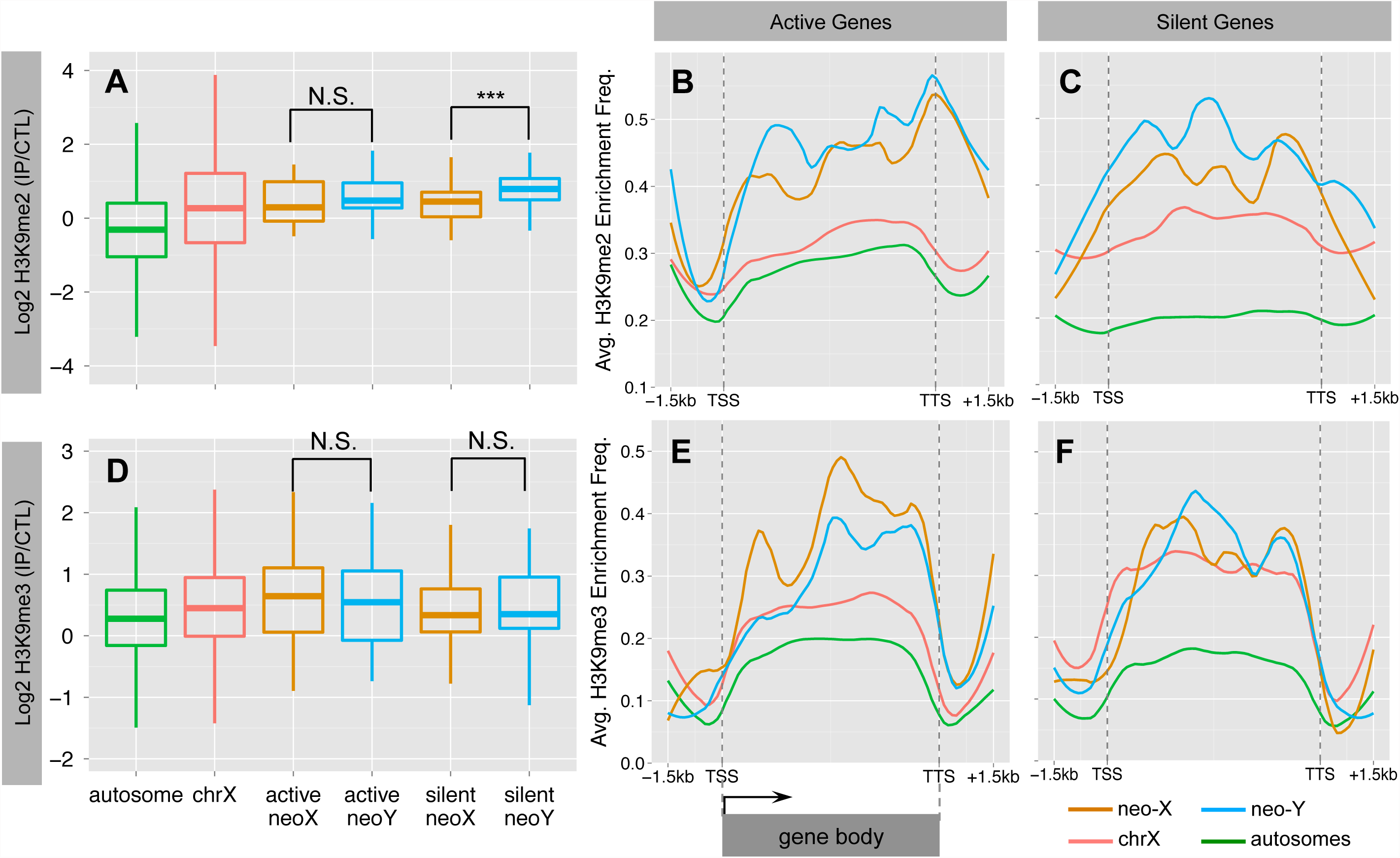
Heterochromatin evolution on the *D. busckii* neo-Y. Shown is the normalized log2 enrichment level of H3K9me2 (**A-C**) or H3K9me3 (**D-F**) over genes on different chromosomes. **A**. Enrichment level of H3K9me2 at silent neo-Y linked genes (in blue) is significantly higher than that of the neo-X (in orange, Wilcoxon test significance level, *P*<0.001:***), chrX (red) and autosomes (green). **B-C.** ‘Metagene’ profiles for H3K9me2 enrichment. Metagene profiles scale all genes of the same chromosome into the same number of bins for calculating average enrichment frequency along the gene body (Methods and Materials). We divide genes into actively transcribed (**B.**) and silent (**C**.) genes based on the gene expression levels of neo-Y alleles. We also include the up- and down-stream 1.5kb flanking regions. **D**. Enrichment level of H3K9me3. **E-F.** Metagene profiles for H3K9me3 enrichment at active (**E**.) and silent (**F**.) genes.

***MSL*-dependent dosage compensation on ancestral X but not on neo-X.**

Most genes on the ancestral X of *D. busckii* are expressed at similar levels in males and females, i.e. they are dosage compensated (**Figure S8**). The molecular mechanism of dosage compensation in *D. busckii* has been unclear, and *in situ* hybridization experiments to polytene chromosomes to stain for components of the *MSL* machinery, using antibodies derived from *D. melanogaster*, have previously failed to identify *MSL* binding on the ancestral X chromosome of *D. busckii* [16]. Instead, an antibody designed against the *POF* protein in *D. melanogaster* was found to coat the entire X chromosome of *D. busckii* in males only [29], and to co-localize with H4K16ac, a histone marker for dosage compensation in Drosophila [46]. This has led to the proposal that *D. busckii* does not utilize the *MSL* machinery to compensate its X chromosome, but instead is using a regulatory mechanism that involves *POF* [47]. However, it is unclear whether the *MSL* antibodies tested are just too diverged to produce a reliable hybridization signal, or if *MSL*-dependent dosage compensation is indeed absent in *D. busckii*.

To evaluate the mechanism of dosage compensation in *D. busckii*, we utilized both bioinformatics and experimental approaches. First, we annotated the intact open reading frames and gene expression patterns of the key *MSL* complex proteins and non-coding RNAs, as well as the *POF* protein and a duplicated copy of *POF* found in *D. busckii*. Transcriptome profiling revealed that *MSL*-2, *POF*, roX-1 and roX-2 non-coding RNA all exhibit male-biased expression patterns (**Figure S9**), similar to their orthologs in *D. melanogaster*. We further performed immunostaining with a new *D. melanogaster MSL-2* antibody, and find weak but male-specific staining of the X chromosome in *D. busckii* (**Figure 5A**). In *D. melanogaster*, the *MSL* complex catalyzes the deposition of the activating histone mark H4K16ac, and ChIP-seq profiling in *D. busckii* clearly reveals that H4K16ac is significantly enriched on the ancestral male X relative to autosomes and the neo-sex chromosomes (Wilcoxon test, *P*<2.2e-16, **Figure 5B**). This is consistent with *MSL*-dependent dosage compensation in *D. busckii*, and orthologous X-linked genes show a significant correlation in their enrichment levels of H4K16ac between larvae samples of *D. busckii* and *D. melanogaster* (Spearman’s rank correlation coefficient: 0.36, *P*<2.2e-16; **Figure 5C**), suggesting that a similar set of genes is being targeted by the dosage compensation complex on the X in both species. Finally, our metagene analysis of the H4K16ac mark reveals a distinctive 3’ bias specifically over active X-linked gene bodies (**Figure 5D**), consistent with the pattern mediated by the *MSL* complex in *D. melanogaster* [46, 48]. Taken together, these results suggest that *D. busckii* shares the same mechanism of dosage compensation for the ancestral X chromosome as *D. melanogaster*, despite their distant phylogenetic relationship (**Figure 2**) and their different sex chromosome karyotype.

**Figure 5.**
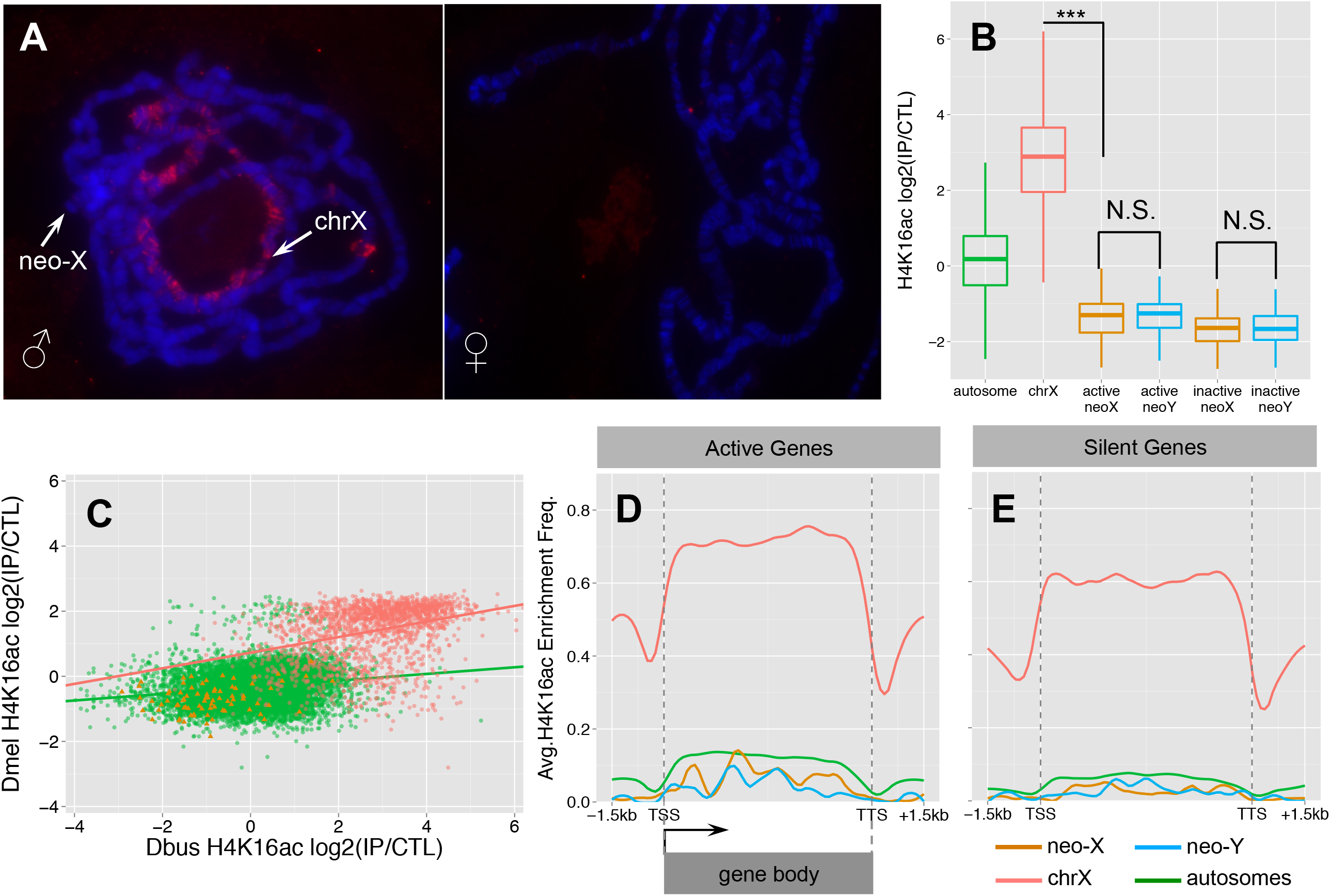
Dosage compensation in male *D. busckii*. **A.** Immunostaining of male and female *D. busckii* polytene chromosomes with *MSL*-2 antibody. The neo-X / X chromosome is marked with an arrow, and the male X shows binding of *MSL*-2 protein. **B.** Comparison of normalized log2 enrichment level of H4K16ac across genes on different chromosomes. Enrichment level of H4K16ac on X-linked genes (red) is significantly higher (Wilcoxon test, *P*<0.001) than on any other chromosomes, while neo-sex linked genes show a significantly lower enrichment level than autosomes (green), and there is no significant difference between the neo-X (orange) and neo-Y (blue) alleles. **C.** Enrichment level of H4K16ac is strongly correlated between orthologous genes of *D. melanogaster* and *D. busckii*. Genes are color-coded according to chromosomal location. **D-E.** Metagene profiles of H4K16ac over active (**D**.) and silent (**E**.) genes. For neo-sex genes, we defined the expression status by the expression level of neo-X alleles. Note that H4K16ac is significantly more enriched at active X-linked genes, and shows a characteristic 3’ binding bias.

Degeneration and down-regulation of neo-Y genes should select for the acquisition of dosage compensation on the *D. busckii* neo-X. If the *MSL*-complex were co-opted on the neo-X in *D. busckii* to achieve dosage compensation, we would expect similar enrichment of H4K16ac along the neo-X in males. Instead, we find that neo-X linked genes are significantly depleted for H4K16ac relative to autosomes (Wilcoxon test, *P*<2.28e-13) (**Figure 5B**), similar to the H4K16ac depletion patterns on the dot in *D. melanogaster* (**Figure S10**). This indicates a lack of *MSL*-dependent dosage compensation on the *D. busckii* neo-X chromosome, in contrast to other neo-sex chromosome systems where a substantial fraction of the neo-Y has become pseudogenized [3, 16]. Instead, it suggests that an ancestrally repressive chromatin structure, as is the case for the dot, may severely constrain the evolution of hyperactive chromatin, despite rampant Y degeneration.

## Discussion

We have performed a detailed investigation of the genomic and epigenomic evolution of the young neo-sex chromosomes of *D. busckii*. All previously studied neo-sex chromosome systems are derived from euchromatic autosomes, but the *D. busckii* neo-sex chromosome originated from the dot chromosome and its unique, more heterochromatic conformation is probably dictating its unusual patterns of chromatin evolution. We found that both the neo-X and neo-Y chromosome are enriched for both H3K9me2/3 relative to other chromosomes, but only H3K9me2 was reported to have a silencing function on the heterochromatic dot chromosome in *D. melanogaster* [45]. Consistent with the idea that increased heterochromatin formation may contribute to the observed down-regulation of neo-Y gene expression (**Figure 3B**), we find that H3K9me2 is enriched at silenced neo-Y linked genes relative to their neo-X homologs, and these genes also tend to become pseudogenized more quickly on the neo-Y. This is consistent with our results in *D. miranda,* and suggests that genes on the neo-Y under lower selective constraints are more likely to become heterochromatic and non-functional early on [3, 44]. In contrast, the H3K9me3 mark is not associated with silent chromatin on the dot chromosome of *D. melanogaster*, and instead enriched along actively transcribed genes on the dot chromosome [45]. We found that neo-Y linked genes show a similar level of H3K9me3 enrichment relative to their neo-X homologs, and no difference between active and silenced genes, suggesting that H3K9me3 does not contribute significantly to expression differences between the neo-X and neo-Y of *D. busckii*. One important caveat in the above analysis is that we can only measure relative expression or histone modification changes on the neo-sex chromosomes, but cannot distinguish whether those changes occurred on the neo-X or neo-Y. It is formally possible that the neo-X has evolved reduced levels of H3K9me2 (but not H3K9me3), relative to the neo-Y. No close relatives of *D. busckii* that lack the neo-sex chromosome fusion are known, preventing us from directly distinguishing between those possibilities.

Two chromosome-wide regulatory systems have been characterized in *D. melanogaster*: one that is mediated by the *MSL* complex and that targets the male X chromosome; and the other that is mediated by *POF* and that targets the dot chromosome in both sexes. *POF* has been shown to bind the nascent RNA of actively transcribed genes on the dot chromosome, and increases levels of expression of these genes [49]. Since some studies placed *D. busckii* as a sister to all other Drosophila species, an apparent lack of *MSL*-binding to the X chromosome [29] has led to the intriguing hypothesis that *POF* may represent an ancestral dosage compensation system. However, our phylogenomic analysis demonstrates that *D. busckii* in fact belongs to the *Drosophila* subgenus (**Figure 2**), and we show that *MSL*-dependent dosage compensation appears to be conserved in *D. busckii*. The *MSL* complex is present in *D. busckii* males and its components show similar male-biased expression patterns as found in *D. melanogaster* (**Figure S9**), it binds the X chromosome of *D. busckii* males (**Figure 5A**), and the H4K16ac dosage compensation mark is found along actively transcribed X-genes in *D. busckii* (**Figure 5B**). This calls for a re-examination of the proposed role of *POF* in dosage compensation in *D. busckii* [29]. Despite clear evidence for dosage compensation of the ancestral X chromosome of *D. busckii* by the *MSL* complex, we found no signs of *MSL*-mediated dosage compensation on its neo-X. Rampant neo-Y degeneration (i.e. almost 60% of neo-Y genes have frameshift mutations or stop codons) should in principle select for the evolution of dosage compensation on the neo-X of *D. busckii*. Indeed, in other Drosophila species with neo-sex chromosomes, dosage compensation was found to evolve rapidly after their formation and degeneration of neo-Y genes, by co-opting the ancestral *MSL* machinery. In *D. miranda*, the neo-X chromosome has evolved partial dosage compensation through the acquisition of novel *MSL*-binding sites that recruit the *MSL*-complex to the neo-X [14, 15], and *MSL*-binding was found to be associated with H4K16ac enrichment. Even older neo-X chromosomes, like the one shared by members of the *D. pseudoobscura* subgroup, have evolved full *MSL*-mediated dosage compensation [3, 16]. In contrast, we did not detect any enrichment of the H4K16ac modification on the neo-X of *D. busckii*. This is probably due to the younger age of the *D. busckii* neo-X chromosome, the fact that the dot chromosome contains only few genes and flies with a single copy of the dot chromosome are fully viable in *D. melanogaster* (due to compensation mediated by POF [50]), and/or the difficulty of evolving a hyper-active chromatin structure for dosage compensation from an ancestrally more heterochromatic background. Our previous work in *D. miranda* showed that dosage compensation preferentially evolves in chromatin regions that are ancestrally active [3], probably due to an antagonism between forming repressive, condensed heterochromatin and hyperactive, open chromatin resulting in dosage compensation [51]. Despite down-regulation of neo-Y genes and a lack of *MSL*-mediated dosage compensation of neo-X genes, we find that transcription of the single neo-X chromosome in males is not simply half that in females, and neo-sex linked genes do not exhibit strong sex-biased expression patterns. This suggests that the down-regulation of neo-Y linked genes is either at least partially compensated by transcriptional buffering mechanism [47], which may play an important role during early sex chromosome differentiation, before the establishment of global dosage compensation on young X chromosomes. Alternatively, a *POF*-mediated regulatory mechanism might compensate for reduced gene dose of neo-Y linked genes. It will be of great interest to further investigate the evolutionary and functional relationship between these two chromosome-wide compensatory mechanisms that have been described in Drosophila.

## Methods

### Data collection genomic DNA and transcriptome

An iso-female line of *D. busckii* provided by J. Larsson and originally caught in Tallinn, Estonia in the year 2000 was used for this study. About 50 virgin adult male and female were used for genomic DNA extraction using Puregene Core Kit A (Qiagen, Inc). Total RNA from about 50 larvae and virgin adult flies of each sex were extracted by RNAeasy Mini Kit (Qiagen, Inc). Library preparation and genomic or poly-A selected transcriptome sequencing were then performed at Beijing Genomic Institute or UC Berkeley Sequencing facility following the standard Illumina protocol. We sequenced the libraries by paired-end sequencing with 90bp read length for all the RNA-seq libraries and most of the genomic libraries, and 50bp for long-insert libraries. Female DNA was sequenced to very high coverage (172 fold, **Table S1**) for *de novo* assembly of a reference genome, and male DNA was sequenced to medium coverage (27 fold).

### Genome assembly & annotation

We assembled the reference genome by ALLPATHS-LG [52] with standard parameters. The output scaffold sequences were aligned to *D. melanogaster* chromosomal sequences (v5.46) downloaded from FlyBase by LASTZ (http://www.bx.psu.edu/∼rsharris/lastz/) using a nucleotide matrix for distant species comparison. Alignment results were filtered using a cutoff of at least 30% of the entire scaffold aligned with 50% sequence identity. We wrote customized perl scripts to build pseudo-chromosomal sequences of *D. busckii*. We further used RepeatMasker and RepeatModeler (<u>http://www.repeatmasker.org</u>) to annotate the repeat content of the *D. busckii* genome.

RNA-seq reads from both sexes were separately aligned to the chromosome sequences of *D. busckii* by tophat [53]. The alignments were then provided to cufflinks [54] for transcriptome annotation. We integrated the annotation results from cufflinks and used a non-redundant protein sequence set of *D. melanogaster* (v5.46) to annotate the *D. busckii* genome using the MAKER pipeline [55]. We annotated 79 out of 80 dot-linked *D. melanogaster* protein-coding genes in the *D. busckii* genome. 71 of them are located on the dot chromosome of *D. busckii,* and the remaining 8 genes are located on the X chromosome or other autosomes, including 4 genes whose *D. virilis* orthologs also map to other chromosomes [56]. The additional 15 genes annotated on the *D. busckii* dot chromosome are either predicted by a combination of RNA-seq evidence and *de novo* open reading frame annotation, or have a *D. melanogaster* ortholog located on another chromosome. We compared the distributions of normalized expression level (measured by Reads Per Kilobase per Million, RPKM) in gene regions and intergenic regions, and used the value where the two distributions separate as a cutoff (log_10_ RPKM =0.65; **Figure S1**) to define genes that are transcriptionally active or not. We analyzed the codon usage bias of all annotated *D. busckii* genes by CodonW (<u>http://codonw.sourceforge.net/</u>).

## SNP calling and allelic specific analyses

We used the standard GATK pipeline [57] for calling SNPs in male and female DNA samples. In brief, sequencing reads were aligned to the *D. busckii* genome with bowtie2 [58] and PCR duplicate reads were removed using the Picard tool (<u>http://broadinstittute.github.io/picard</u>). We used UnifiedGenotyper for calling variants, and discarded SNPs/indels with low qualities (Quality<30), low coverage (Depth<5), strand biases or clustering patterns for initial SNP filtering. To account for the different sequencing coverage of the male and female samples, we further plot the distributions of variant qualities of male and female SNPs to determine a different variant quality cutoff for the second round of filtering. We identified a total of 16977 heterozygous SNP sites from the male sample and only 496 female heterozygous sites on the dot chromosome. After excluding the sites that are shared by both sexes, we used the quality-filtered male-specific SNPs/indels as the putative fixed neo-X/Y divergence sites, and introduced the alternative nucleotides to the reference neo-X genome to produce the reference genomic sequence of the neo-Y chromosome. Note that only individuals from a single inbred line were sequenced; this means that some of the fixed differences between the neo-X and neo-Y are not actually fixed in the population but may be segregating on either chromosome. Based on the female-specific heterozygous sites, we estimated that only 1.5% of the divergence sites maybe derived from segregating polymorphic sites. We then used GeneWise [59] and annotated the non-functional genes of the neo-Y using the proteins annotated from the female reference genome as query.

To analyze neo-X and neo-Y allele-specific gene expression and histone profiles (see below), we aligned the male RNA-seq or ChIP-seq reads against the female reference genome and specifically collected reads that overlapped the male-specific SNP sites. These reads encompass informative neo-X/Y divergence sites, and we used customized perl scripts to assign their linkage to either the neo-X or neo-Y, dependent on whether the SNP is male-specific or not. To correct for potential mapping biases, we normalized the count of RNA-seq reads against the DNA-seq reads from males, whose ratios between neo-X and neo-Y alleles are expected to be 1. To test the significance of biased gene expression between neo-X/Y alleles, we used Fisher’s tests with the allelic-specific DNA-seq read count and allelic-specific RNA-seq read count of the neo-X or neo-Y allele for the 2×2 table. This should account for potential mapping biases of neo-X and neo-Y derived reads, and their ratio is expected to be similar between neo-X/Y alleles if they are transcribing at a similar level. Since the enrichment of ChIP-seq profiles is calculated by normalizing against the input DNA-seq control, we did not do any further correction. When comparing the binding level between the neo-X/Y alleles or different chromosomes, we calculated the ratio of aligned read numbers of ChIP experiment vs. input DNA control, spanning the gene body and 1.5 kb flanking regions.

## Phylogenomic and PAML analyses

We collected CDS sequences from *D. pseudoobscura* (v3.1), *D. virilis* (v1.2), *D. willistoni* (v1.3), *D. grimshawi* (v1.3) and *D. albomicans* from FlyBase, and two Diptera species *Scaptodrosophila lebanonensis* and zoophilic fruitfly (*Phortica variegata*) whose genomes have been recently produced in our lab [19, 36]. Orthologous relationships of genes between species were determined through reciprocal BLAST or precomputed annotation from FlyBase. We aligned all the orthologous sequences for the same gene by translatorX [60], a program that performs codon-based nucleotide sequence alignment and removed low-quality alignment regions by Gblock [61]. The alignments were then concatenated and provided to RAxML for constructing maximum-likelihood trees with the GTRCAT algorithm, with *P. variegate* assigned as an outgroup to all Drosophila species. We bootstrapped the tree 1,000 times and calculated confidence values for each node as described in the manual of RAxML [62].

Branch-specific evolutionary rates were calculated for the resulting high-confidence tree using the PAML package [63]. To collect data for as many genes as possible, we only used *D. melanogaster* and *D. virilis* as outgroups of *D. busckii* in the input tree. We calculated lineage specific synonymous or nonsynonymous substitution rates using codeml under the ‘free-ratio’ model, which assumes each phylogenetic branch has a different rate of evolution.

## Polytene chromosome staining and ChIP-seq

Polytene chromosomes were dissected from male third instar larvae and processed for immunostaining with primary *MSL*-2 antibody (Santa Cruz Biotechnology, sc-32458, dilution ratio: 1:10, room temperature, overnight) and secondary fluorescence antibody Alexa Fluor 555 Dye (Life Technologies, room temperature, 2 hours). Approximately 5g of male third instar larvae were used for chromatin extraction. Chromatin was cross-linked with formaldehyde and sheared by sonication. Chromatin pull-down with IgG agarose beads (Sigma, A2909) was performed as described previously [64]. We used the following antibodies for ChIP-seq experiments: (1) H3K9me3 (Abcam ab8898; 3 μl/IP) (2) anti-H4K16ac (Millipore 07-329; 5 μl/IP) (3) H3K9me2 (Abcam ab1220; 3μl/IP). Immunoprecipitated and input DNAs were purified and processed according to the standard paired-end Solexa library preparation protocol. Paired-end 100-bp DNA sequencing was performed on the Illumina Genome Analyzer located at UC Berkeley Vincent J. Coates Genomic Sequencing Facility. ChIP-seq and input control reads were aligned to the *D. busckii* genome by bowtie2 [58]. The resulting alignments were filtered using a cutoff for mapping quality higher than 30, and provided to MACS [65] to call peaks of enrichment along the chromosomes. We use MEME [66] to identify targeting sequence motifs within peak regions. For metagene analyses, we first determine a cutoff to define ‘bound’ or ‘unbound’ states of certain chromatin marks within each scaled bin of genes or flanking regions, by comparing the distribution of their normalized enrichment levels between chromosomes (**Figure S11**). Then for each bin, we calculated the average bound level across all the studied genes, after dividing them into different groups of chromosomes and active/silent genes.

ChIP-seq data of male *D. melanogaster* is downloaded from NCBI SRA database (accession#: PRJEB3015) [67] and orthologous relationship between *D. busckii* and *D. melanogaster* genes was determined using reciprocally best BLAST searches.

## Acknowledgments

We thank Jan Larsson and Zaak Walton for technical assistance, and Nicolas Gompel at LMU Munich for taking the amazing Drosophila photos. Reads, genome assembly and annotation generated in this project are all deposited under the Bioproject Number PRJNA274996 on NCBI. This project is supported by NIH Grants (R01GM076007, R01GM101255 and R01GM093182) to D.B.

## Supplementary Figures

**Figure S1.** Distribution of gene expression levels on different chromosomes

Shown are density plots of gene expression for protein coding genes on autosomes (green), the X chromosome (red) and dot chromosome (orange) of *D. busckii* male larvae and adults. We also plot the expression level of intergenic regions (dotted line), to determine a cutoff value (dashed line) for defining actively transcribed genes.

**Figure S2.** Male and female coverage patterns of neo-sex linked genes

Shown is log10 based read coverage of neo-sex chromosome genes in males (x-axis) and females (y-axis). A similar level of coverage between sexes indicates that none of the neo-Y genes are deleted.

**Figure S3.** Codon usage bias pattern across different chromosomes

We compare levels of codon usage bias between genes on different chromosomes, using the neo-X sequences for the dot chromosome. Different measurements of codon usage bias, including codon bias index (CBI), frequency of optimal codons (as defined by *D. melanogaster*, Fop) and codon adaptation index (CAI) consistently show that dot-linked genes have reduced levels of codon usage bias.

**Figure S4.** Comparisons of rates of protein evolution measured by the nonsynonymous substitution rate (*K*_a_), synonymous substitution rate (*K*_s_), and their ratio (*K*_a_/*K*_s_) for different chromosomes

Shown are boxplots for *K*_a_, *K*_s_ and the *K*_a_/*K*_s_ ratio, for genes linked to autosomes, the X chromosome, and ancestral *K*_a_, *K*_s_ and *K*_a_/*K*_s_ ratios before neo-sex divergence for putatively functional neo-Y linked genes (func) and non-functional neo-Y linked genes (psd). We show Wilcoxon test significance level: *P* < 0.05: *, *P*<0.01: **, *P*<0.001: ***.

**Figure S5.** *D. busckii* gene expression in male larvae

**A.** Shown is the relative male larvae expression of neo-X vs. neo-Y along the neo-sex chromosomes, with functional genes in green and pseudogenes in red. **B.** Boxplots of gene expression level of different chromosomes, with neo-sex linked genes divided into functional (func) and non-functional (psd) neo-Y genes, and their corresponding neo-X homologs (diploid vs. hemizygous neo-Xs, dpd vs. hmz). We show Wilcoxon test significance level: *P* < 0.05: *, *P*<0.01: **, *P*<0.001: ***.

**Figure S6.** Comparing enrichment levels of heterochromatin marks between functional and non-functional neo-Y genes

Boxplots showing the H3K9me2/3 enrichment level of functional and nonfunctional neo-Y linked genes (in blue), and their corresponding neo-X homologs. H3K9me2 but not H3K9me3 is significantly (Wilcoxon test, *P*<0.05) enriched on the non-functional neo-Y genes relative to their neo-X homologs.

**Figure S7.** -Y H3K9me2 enrichment level at neo-Y genes is negatively correlated with their expression level

Shown are the normalized enrichment levels of H3K9me2 and H3K9me3 for neo-sex linked genes vs. their allelic gene expression level. Functional neo-Y genes and their neo-X homologs are in green, and non-functional neo-Y genes and their neo-X homologs in red. Only H3K9me2 shows a significant negative correlation (*F*-statistic test, *P*<0.05) with gene expression level on the neo-Y. Note that non-functional neo-Y genes show a stronger negative correlation between H3K9me2 enrichment and expression level than functional neo-Y genes.

**Figure S8.** Distribution of male vs. female expression ratios of *D. busckii* genes

Density plot of male vs. female adult gene expression ratio, with X-linked genes in red, and autosomal genes in green. Most genes show equal expression levels between sexes, resulting in a peak centered at 0. Due to the demasculinization of X-linked genes (**Figure S1**), this peak is shifted from 0 toward a lower relative expression in males.

**Figure S9.** Gene expression patterns of *MSL* complex proteins, noncoding RNAs and *POF* proteins

Shown are Gbrowser plots of *MSL* complex proteins, roX non-coding RNAs, and *POF* protein and *POF* duplicate protein of *D. busckii.* Their sex-biased gene expression pattern is consistent with their *D. melanogaster* orthologs.

**Figure S10.** Normalized H4K16ac enrichment of genes on different *D. melanogaster* chromosomes

Shown are boxplots of log2 normalized H4K16ac enrichment levels from salivary glands of third instar male *D. melanogaster* larvae [67]. Note that the dot chromosome is deficient for the active H4K16ac mark.

**Figure S11** Distribution of enrichment levels of chromatin marks across different chromosomes

Shown are histograms of log2 normalized enrichment level of different chromatin marks within scaled bins of genes on different chromosomes from third instar larvae of male *D. busckii*. We determine an arbitrary cutoff (the dashed line) to define ‘bound’ or ‘unbound’ genes for a certain mark, which separates the distribution of sex or the dot chromosome from others. The chromosomes are named after their homologous *D. melanogaster* chromosomes.

**Table S1** Sequencing coverage of the *D. busckii* genome

## References

1. Bull JJ. Evolution of sex determining mechanisms: The Benjamin/Cummings Publishing Company, Inc.; 1983.

2. Bachtrog D. Y-chromosome evolution: emerging insights into processes of Y-chromosome degeneration. Nat Rev Genet. 2013;14(2):113–24. DOI: Doi 10.1038/Nrg3366. PubMed PMID: WOS:000314622000011.

3. Zhou Q, Ellison CE, Kaiser VB, Alekseyenko AA, Gorchakov AA, Bachtrog D. The epigenome of evolving Drosophila neo-sex chromosomes: dosage compensation and heterochromatin formation. PLoS biology. 2013;11(11):e1001711. Epub 2013/11/23. DOI: 10.1371/journal.pbio.1001711. PubMed PMID: 24265597; PubMed Central PMCID: PMC3825665.

4. Conrad T, Akhtar A. Dosage compensation in Drosophila melanogaster: epigenetic fine-tuning of chromosome-wide transcription. Nat Rev Genet. 2011;13(2):123–34. DOI: 10.1038/nrg3124. PubMed PMID: 22251873.

5. Disteche CM. Dosage compensation of the sex chromosomes. Annual review of genetics. 2012; 46:537–60. DOI: 10.1146/annurev-genet-110711-155454. PubMed PMID: 22974302; PubMed Central PMCID: PMC3767307.

6. Marais GA, Nicolas M, Bergero R, Chambrier P, Kejnovsky E, Moneger F, et al. Evidence for degeneration of the Y chromosome in the dioecious plant Silene latifolia. Current biology : CB. 2008;18(7):545–9. DOI: 10.1016/j.cub.2008.03.023. PubMed PMID: 18394889.

7. Zhang W, Wang X, Yu Q, Ming R, Jiang J. DNA methylation and heterochromatinization in the male-specific region of the primitive Y chromosome of papaya. Genome research. 2008;18(12):1938–43. DOI: 10.1101/gr.078808.108. PubMed PMID: 18593814; PubMed Central PMCID: PMC2593574.

8. Yoshida K, Makino T, Yamaguchi K, Shigenobu S, Hasebe M, Kawata M, et al. Sex chromosome turnover contributes to genomic divergence between incipient stickleback species. PLoS Genet. 2014;10(3):e1004223. DOI: 10.1371/journal.pgen.1004223.

9. Zhou Q, Bachtrog D. Chromosome-wide gene silencing initiates Y degeneration in Drosophila. Current biology : CB. 2012;22(6):522–5. DOI: 10.1016/j.cub.2012.01.057. PubMed PMID: 22365853.

10. Charlesworth B, Charlesworth D. The degeneration of Y chromosomes. Philos Trans R Soc Lond B Biol Sci. 2000;355(1403):1563–72. Epub 2000/12/29. PubMed PMID: 11127901; PubMed Central PMCID: PMC1692900.

11. Zhou Q, Bachtrog D. Sex-specific adaptation drives early sex chromosome evolution in Drosophila. Science. 2012;337(6092):341–5. Epub 2012/07/24. DOI: 10.1126/science.1225385. PubMed PMID: 22822149.

12. Steinemann M, Steinemann S. Enigma of Y chromosome degeneration: neo-Y and neo-X chromosomes of Drosophila miranda a model for sex chromosome evolution. Genetica. 1998; 102-103(1-6):409–20. Epub 1998/08/28. PubMed PMID: 9720292.

13. Bachtrog D, Hom E, Wong KM, Maside X, de Jong P. Genomic degradation of a young Y chromosome in Drosophila miranda. Genome Biol. 2008;9(2):R30. Epub 2008/02/14. DOI: gb-2008-9-2-r30 [pii] 10.1186/gb-2008-9-2-r30. PubMed PMID: 18269752.

14. Alekseyenko AA, Ellison CE, Gorchakov AA, Zhou Q, Kaiser VB, Toda N, et al. Conservation and de novo acquisition of dosage compensation on newly evolved sex chromosomes in Drosophila. Genes & development. 2013;27(8):853–8. Epub 2013/05/01. DOI: 10.1101/gad.215426.113. PubMed PMID: 23630075; PubMed Central PMCID: PMC3650223.

15. Ellison CE, Bachtrog D. Dosage compensation via transposable element mediated rewiring of a regulatory network. Science. 2013;342(6160):846–50. Epub 2013/11/16. DOI: 10.1126/science.1239552. PubMed PMID: 24233721.

16. Marin I, Franke A, Bashaw GJ, Baker BS. The dosage compensation system of Drosophila is co-opted by newly evolved X chromosomes. Nature. 1996;383(6596):160–3. DOI: Doi 10.1038/383160a0. PubMed PMID: WOS:A1996VG14800049.

17. Krivshenko J. New Evidence for the Homology of the Short Euchromatic Elements of the X and Y Chromosomes of Drosophila Busckii with the Microchromosome of Drosophila Melanogaster. Genetics. 1959;44(6):1027–40. Epub 1959/11/01. PubMed PMID: 17247874; PubMed Central PMCID: PMC1224414.

18. Krivshenko JD. A cytogenetic study of the X chromosome of Drosophila busckii and its relation to phylogeny. Proceedings of the National Academy of Sciences of the United States of America. 1955;41(12):1071–9. Epub 1955/12/15. PubMed PMID: 16589798; PubMed Central PMCID: PMC528199.

19. Vicoso B, Bachtrog D. Reversal of an ancient sex chromosome to an autosome in Drosophila. Nature. 2013;499(7458):332–5. Epub 2013/06/25. DOI: 10.1038/nature12235. PubMed PMID: 23792562.

20. Vicoso B, Bachtrog D. Numerous transitions of sex chromosomes in Diptera. PLoS Biology. 2015;13(4):e1002078.

21. Hochman B. The fourth chromosome of Drosophila melanogaster. Genetic Biology of the Drosophila (USA). 1976.

22. Jensen MA, Charlesworth B, Kreitman M. Patterns of genetic variation at a chromosome 4 locus of Drosophila melanogaster and D-simulans. Genetics. 2002;160(2):493–507. PubMed PMID: WOS:000174097600015.

23. Arguello JR, Zhang Y, Kado T, Fan C, Zhao R, Innan H, et al. Recombination yet inefficient selection along the Drosophila melanogaster subgroup’s fourth chromosome. Molecular biology and evolution. 2010;27(4):848–61. DOI: 10.1093/molbev/msp291. PubMed PMID: 20008457; PubMed Central PMCID: PMC2877538.

24. Adams MD, Celniker SE, Holt RA, Evans CA, Gocayne JD, Amanatides PG, et al. The genome sequence of Drosophila melanogaster. Science. 2000;287(5461):2185–95. Epub 2000/03/25. PubMed PMID: 10731132.

25. Haddrill PR, Halligan DL, Tomaras D, Charlesworth B. Reduced efficacy of selection in regions of the Drosophila genome that lack crossing over. Genome biology. 2007;8(2):R18.

26. Campos JL, Charlesworth B, Haddrill PR. Molecular evolution in nonrecombining regions of the Drosophila melanogaster genome. Genome biology and evolution. 2012;4(3):278–88. DOI: 10.1093/gbe/evs010. PubMed PMID: 22275518; PubMed Central PMCID: PMC3318434.

27. Riddle NC, Minoda A, Kharchenko PV, Alekseyenko AA, Schwartz YB, Tolstorukov MY, et al. Plasticity in patterns of histone modifications and chromosomal proteins in Drosophila heterochromatin. Genome research. 2011;21(2):147–63. Epub 2010/12/24. DOI: 10.1101/gr.110098.110. PubMed PMID: 21177972; PubMed Central PMCID: PMC3032919.

28. Larsson J, Chen JD, Rasheva V, Rasmuson-Lestander A, Pirrotta V. Painting of fourth, a chromosome-specific protein in Drosophila. Proceedings of the National Academy of Sciences of the United States of America. 2001;98(11):6273–8. Epub 2001/05/17. DOI: 10.1073/pnas.111581298. PubMed PMID: 11353870; PubMed Central PMCID: PMC33458.

29. Larsson J, Svensson MJ, Stenberg P, Makitalo M. Painting of fourth in genus Drosophila suggests autosome-specific gene regulation. Proceedings of the National Academy of Sciences of the United States of America. 2004;101(26):9728–33. Epub 2004/06/24. DOI: 10.1073/pnas.0400978101. PubMed PMID: 15210994; PubMed Central PMCID: PMC470743.

30. Grimaldi DA. A phylogenetic, revised classification of genera in the Drosophilidae (Diptera). B Am Mus Nat Hist. 1990;(197):1–139. PubMed PMID: WOS:A1990EF89200001.

31. Larsson J, Meller VH. Dosage compensation, the origin and the afterlife of sex chromosomes. Chromosome research : an international journal on the molecular, supramolecular and evolutionary aspects of chromosome biology. 2006;14(4):417–31. Epub 2006/07/06. DOI: 10.1007/s10577-006-1064-3. PubMed PMID: 16821137.

32. Bhutkar A, Schaeffer SW, Russo SM, Xu M, Smith TE, Gelbart WM. Chromosomal rearrangement inferred from comparisons of 12 Drosophila genomes. Genetics. 2008;179(3):1657–80. DOI: Doi 10.1534/Genetics.107.086108. PubMed PMID: WOS:000258313400039.

33. Graveley BR, Brooks AN, Carlson JW, Duff MO, Landolin JM, Yang L, et al. The developmental transcriptome of Drosophila melanogaster. Nature. 2011;471(7339):473–9. Epub 2010/12/24. DOI: 10.1038/nature09715. PubMed PMID: 21179090; PubMed Central PMCID: PMC3075879.

34. Sturgill D, Zhang Y, Parisi M, Oliver B. Demasculinization of X chromosomes in the Drosophila genus. Nature. 2007;450(7167):238–41. DOI: 10.1038/nature06330. PubMed PMID: 17994090; PubMed Central PMCID: PMC2386140.

35. van der Linde K, Houle D, Spicer GS, Steppan SJ. A supermatrix-based molecular phylogeny of the family Drosophilidae. Genet Res. 2010;92(1):25–38. Epub 2010/05/04. DOI: 10.1017/S001667231000008X1. PubMed PMID: 20433773.

36. Vicoso B, Bachtrog D. Numerous transitions of sex chromosomes in Diptera. PLoS biology. 2015;13(4):e1002078. DOI: 10.1371/journal.pbio.1002078.

37. Kwiatowski J, Ayala FJ. Phylogeny of Drosophila and related genera: conflict between molecular and anatomical analyses. Mol Phylogenet Evol. 1999;13(2):319–28. DOI: 10.1006/mpev.1999.0657. PubMed PMID: 10603260.

38. Schrider DR, Houle D, Lynch M, Hahn MW. Rates and genomic consequences of spontaneous mutational events in Drosophila melanogaster. Genetics. 2013;194(4):937- DOI: 10.1534/genetics.113.151670. PubMed PMID: 23733788; PubMed Central PMCID: PMC3730921.

39. Bachtrog D. The temporal dynamics of processes underlying Y chromosome degeneration. Genetics. 2008;179(3):1513–25. DOI: 10.1534/genetics.107.084012. PubMed PMID: 18562655; PubMed Central PMCID: PMC2475751.

40. Kaiser VB, Charlesworth B. The effects of deleterious mutations on evolution in non-recombining genomes. Trends Genet. 2009;25(1):9–12. DOI: 10.1016/j.tig.2008.10.009. PubMed PMID: 19027982.

41. Haddrill PR, Halligan DL, Tomaras D, Charlesworth B. Reduced efficacy of selection in regions of the Drosophila genome that lack crossing over. Genome biology. 2007;8(2). DOI: Artn R18 Doi 10.1186/Gb-2007-8-2-R18. PubMed PMID: WOS:000246076300006.

42. Lindsley DL, Sandler L, Baker BS, Carpenter AT, Denell RE, Hall JC, et al. Segmental aneuploidy and the genetic gross structure of the Drosophila genome. Genetics. 1972;71(1):157–84. Epub 1972/05/01. PubMed PMID: 4624779; PubMed Central PMCID: PMC1212769.

43. Powell JR, Dion K, Papaceit M, Aguade M, Vicario S, Garrick RC. Nonrecombining genes in a recombination environment: the Drosophila “dot” chromosome. Molecular biology and evolution. 2011;28(1):825–33. DOI: 10.1093/molbev/msq258. PubMed PMID: 20940345; PubMed Central PMCID: PMC3002241.

44. Kaiser VB, Zhou Q, Bachtrog D. Nonrandom gene loss from the Drosophila miranda neo-Y chromosome. Genome biology and evolution. 2011;3:1329–37. Epub 2011/10/12. DOI: 10.1093/gbe/evr103. PubMed PMID: 21987387; PubMed Central PMCID: PMC3236567.

45. Riddle NC, Jung YL, Gu TT, Alekseyenko AA, Asker D, Gui HX, et al. Enrichment of HP1a on Drosophila Chromosome 4 Genes Creates an Alternate Chromatin Structure Critical for Regulation in this Heterochromatic Domain. PLoS Genet. 2012; 8(9). DOI: Artn E1002954 Doi 10.1371/Journal.Pgen.1002954. PubMed PMID: WOS:000309817900029.

46. Gelbart ME, Larschan E, Peng S, Park PJ, Kuroda MI. Drosophila MSL complex globally acetylates H4K16 on the male X chromosome for dosage compensation. Nature structural & molecular biology. 2009;16(8):825–32. Epub 2009/08/04. DOI: 10.1038/nsmb.1644. PubMed PMID: 19648925; PubMed Central PMCID: PMC2722042.

47. Stenberg P, Larsson J. Buffering and the evolution of chromosome-wide gene regulation. Chromosoma. 2011;120(3):213–25. Epub 2011/04/21. DOI: 10.1007/s00412-011-0319-8. PubMed PMID: 21505791; PubMed Central PMCID: PMC3098985.

48. Kind J, Vaquerizas JM, Gebhardt P, Gentzel M, Luscombe NM, Bertone P, et al. Genome-wide analysis reveals MOF as a key regulator of dosage compensation and gene expression in Drosophila. Cell. 2008;133(5):813–28.

49. Johansson AM, Stenberg P, Allgardsson A, Larsson J. POF regulates the expression of genes on the fourth chromosome in Drosophila melanogaster by binding to nascent RNA. Mol Cell Biol. 2012;32(11):2121–34. DOI: 10.1128/MCB.06622-11. PubMed PMID: 22473994; PubMed Central PMCID: PMC3372238.

50. Johansson AM, Stenberg P, Bernhardsson C, Larsson J. Painting of fourth and chromosome-wide regulation of the 4th chromosome in Drosophila melanogaster. EMBO J. 2007;26: 2307–2316.

51. Spierer A, Begeot F, Spierer P, Delattre M. SU(VAR)3-7 links heterochromatin and dosage compensation in Drosophila. PLoS Genet. 2008;4(5):e1000066. Epub 2008/05/03. DOI: 10.1371/journal.pgen.1000066. PubMed PMID: 18451980; PubMed Central PMCID: PMC2320979.

52. Gnerre S, Maccallum I, Przybylski D, Ribeiro FJ, Burton JN, Walker BJ, et al. High-quality draft assemblies of mammalian genomes from massively parallel sequence data. Proceedings of the National Academy of Sciences of the United States of America.2011;108(4):1513–8. Epub 2010/12/29. DOI: 10.1073/pnas.1017351108. PubMed PMID:21187386; PubMed Central PMCID: PMC3029755.

53. Trapnell C, Pachter L, Salzberg SL. TopHat: discovering splice junctions with RNA-Seq. Bioinformatics. 2009;25(9):1105–11. Epub 2009/03/18. DOI: 10.1093/bioinformatics/btp120. PubMed PMID: 19289445; PubMed Central PMCID: PMC2672628.

54. Trapnell C, Williams BA, Pertea G, Mortazavi A, Kwan G, van Baren MJ, et al. Transcript assembly and quantification by RNA-Seq reveals unannotated transcripts and isoform switching during cell differentiation. Nature biotechnology. 2010;28(5):511–5. Epub 2010/05/04. DOI: 10.1038/nbt.1621. PubMed PMID: 20436464; PubMed Central PMCID: PMC3146043.

55. Cantarel BL, Korf I, Robb SM, Parra G, Ross E, Moore B, et al. MAKER: an easy-to-use annotation pipeline designed for emerging model organism genomes. Genome research. 2008;18(1):188–96. Epub 2007/11/21. DOI: 10.1101/gr.6743907. PubMed PMID: 18025269; PubMed Central PMCID: PMC2134774.

56. Leung W, Shaffer CD, Cordonnier T, Wong J, Itano MS, Slawson Tempel EE, et al. Evolution of a distinct genomic domain in Drosophila: comparative analysis of the dot chromosome in Drosophila melanogaster and Drosophila virilis. Genetics. 2010;185(4):1519–34. DOI: 10.1534/genetics.110.116129. PubMed PMID: 20479145; PubMed Central PMCID: PMC2927774.

57. DePristo MA, Banks E, Poplin R, Garimella KV, Maguire JR, Hartl C, et al. A framework for variation discovery and genotyping using next-generation DNA sequencing data. Nat Genet. 2011;43(5):491-+. DOI: Doi 10.1038/Ng.806. PubMed PMID: WOS:000289972600023.

58. Langmead B, Salzberg SL. Fast gapped-read alignment with Bowtie 2. Nature methods. 2012;9(4):357–9. Epub 2012/03/06. DOI: 10.1038/nmeth.1923. PubMed PMID: 22388286; PubMed Central PMCID: PMC3322381.

59. Birney E, Clamp M, Durbin R. GeneWise and genomewise. Genome Res. 2004;14(5):988–95. DOI: Doi 10.1101/Gr.1865504. PubMed PMID: WOS:000221171700026.

60. Abascal F, Zardoya R, Telford MJ. TranslatorX: multiple alignment of nucleotide sequences guided by amino acid translations. Nucleic Acids research. 2010;38(Web Server issue):W7-13. Epub 2010/05/04. DOI: 10.1093/nar/gkq291. PubMed PMID: 20435676; PubMed Central PMCID: PMC2896173.

61. Castresana J. Selection of conserved blocks from multiple alignments for their use in phylogenetic analysis. Molecular biology and evolution. 2000;17(4):540–52. Epub 2000/03/31. PubMed PMID: 10742046.

62. Stamatakis A. RAxML version 8: a tool for phylogenetic analysis and post-analysis of large phylogenies. Bioinformatics. 2014. Epub 2014/01/24. DOI: 10.1093/bioinformatics/btu033. PubMed PMID: 24451623.

63. Yang Z. PAML 4: phylogenetic analysis by maximum likelihood. Molecular biology and evolution. 2007;24(8):1586–91. Epub 2007/05/08. DOI: 10.1093/molbev/msm088. PubMed PMID: 17483113.

64. Alekseyenko AA, Peng SY, Larschan E, Gorchakov AA, Lee OK, Kharchenko P, et al. A sequence motif within chromatin entry sites directs MSL establishment on the Drosophila X chromosome. Cell. 2008;134(4):599–609. DOI: Doi 10.1016/J.Cell.2008.06.033. PubMed PMID: WOS:000258665800014.

65. Zhang Y, Liu T, Meyer CA, Eeckhoute J, Johnson DS, Bernstein BE, et al. Model-based analysis of ChIP-Seq (MACS). Genome biology. 2008;9(9):R137. Epub 2008/09/19. DOI: 10.1186/gb-2008-9-9-r137. PubMed PMID: 18798982; PubMed Central PMCID: PMC2592715.

66. Machanick P, Bailey TL. MEME-ChIP: motif analysis of large DNA datasets. Bioinformatics. 2011;27(12):1696–7. Epub 2011/04/14. DOI: 10.1093/bioinformatics/btr189. PubMed PMID: 21486936; PubMed Central PMCID: PMC3106185.

67. Conrad T, Cavalli FM, Vaquerizas JM, Luscombe NM, Akhtar A. Drosophila dosage compensation involves enhanced Pol II recruitment to male X-linked promoters. Science. 2012;337(6095):742–6. DOI: 10.1126/science.1221428. PubMed PMID: 22821985.

